# SUN5 forms a regular protein lattice reinforcing the sperm head-tail junction

**DOI:** 10.1101/2025.07.29.667520

**Authors:** Jonas Moecking, Svetlana Doroshev, Miguel R. Leung, Tzviya Zeev-Ben-Mordehai

**Affiliations:** Bijvoet Centre for Biomolecular Research, Utrecht University, 3584 CG Utrecht, The Netherlands; Hubrecht Institute-KNAW & University Medical Center Utrecht, 3584 CT, Utrecht, The Netherlands

**Keywords:** LINC complex, SUN domain, cryo-FIB milling, cryo-electron tomography, subtomogram averaging, sperm

## Abstract

Linker of nucleo- and cytoskeleton (LINC) complexes reside in the nuclear envelope, the double-membrane surrounding the nucleus, where they establish a physical bridge between nucleus and cytoplasm. LINC complexes are conserved throughout the tree of life and are present in most nucleated cell types in the human body. They play a major role in signal transduction across the nuclear envelope and in regulating nuclear morphology. One of the most drastic nuclear remodeling events occurs during sperm maturation. Multiple sperm-specific LINC complexes are essential for the sperm cell to adapt its highly streamlined nuclear shape and to secure a stable connection between sperm head and tail. Importantly, mutations in the LINC protein SUN5 result in head-tail detachment, also referred to as acephalic spermatozoa syndrome, rendering affected individuals infertile. Here, using super-resolution fluorescence microscopy we find that sperm-specific SUN5 localizes to the base of the human sperm head and by applying in situ cryo-electron tomography we find an extensive two-dimensional lattice in the nuclear envelope at this region. Moreover, this lattice appears to maintain a consistent close apposition between the inner and outer nuclear membranes. Further structural analysis supports a model in which SUN5 forms trimers that laterally interact at the outer nuclear membrane. Overall, this study sheds light on nuclear envelope organization in the highly streamlined sperm cell, providing mechanistic insights into uniform nuclear envelope spacing maintained by a LINC lattice and rationalizing disruptive effects of SUN5 mutations on the sperm head-tail junction.

## Introduction

The nucleus is a highly dynamic organelle that can change its size, shape and subcellular position in response to cellular or mechanical cues (1). This nuclear adaptation is important for many cellular processes including mitosis, cell migration and differentiation (1, 2). Changes in nuclear morphology are mainly mediated by the double lipid bilayer enclosing the nucleus, the nuclear envelope (NE), consisting of the inner (INM) and outer (ONM) nuclear membrane (3). INM and ONM harbor a multitude of proteins that facilitate nucleocytoplasmic transport, signal transduction or nuclear reshaping, and defects in NE proteins resulting in dysregulated nuclear morphology are associated with specific diseases (1–4).

Linker of nucleo- and cytoskeleton (LINC) complexes are key proteins of the NE and they are conserved in yeast, flies, fish, and mammals (5, 6). As their name indicates, they establish a direct physical connection between elements of the nucleoskeleton, like the nuclear lamina, and elements of the cytoskeleton, like microtubules (6). Their functions range from regulation of nuclear migration and polarity to mechanotransduction. They are typically composed of a SUN (Sad1 and UNC domain containing) protein and a Nesprin (Nuclear envelope spectrin repeat) protein (5–8). SUN proteins are INM transmembrane proteins with their N-terminus residing in the nucleus and typically binding nucleoskeletal proteins, like Lamins, and their C-terminal SUN domain localizing to the perinuclear space (PNS) (5). In the PNS, the SUN domain interacts with the C-terminus of a Nesprin protein, which resides in the ONM and reaches into the cytoplasm with their N-terminal spectrin repeats to bind cytoskeletal elements like actin and microtubules (5, 6, 9–11). Crystal structures of purified SUN domains from SUN1 and SUN2 revealed homotrimers with each protomer bound to a short C-terminal Nesprin peptide (12–14). The region before the SUN domain forms coiled-coils in SUN1 and SUN2 that act as regulators and are important for Nesprin recruitment (15, 16). Intriguingly, purified SUN domain trimers can also form higher order multimers, including dimers of trimers or trimers of trimers (17–19). Based on these in vitro observations the formation of 2D LINC lattices has been proposed (8, 11–14, 20). Yet, whether and how LINC complexes organize into such higher-order assemblies in the NE was not determined experimentally.

The reduction of the sperm nucleus during maturation is one of the most dramatic nuclear remodeling events in mammals, with the nucleus of a mature sperm cell being only about 1/20th of the size of a somatic cell nucleus (21). The fact that out of the five mammalian SUN proteins SUN3, SUN4 and SUN5 as well as the SUN1η isoform are testis-specific, suggests that LINC complexes have a significant role in adapting the highly streamlined shape of the sperm nucleus (5, 22–28). Indeed, SUN3-5 play critical roles in reshaping and positioning the nucleus during different stages of sperm maturation and in forming a stable connection between sperm head and tail (29, 30). In mice, SUN3 is expressed during the postmeiotic stages of spermatogenesis and interacts with Nesprin1 (22). Immunofluorescence experiments showed that SUN3 localizes to the attachment sites between the NE and the manchette, a transient microtubule-based structure crucial for sperm head formation. SUN4 displays a similar expression and localization profile during early spermatogenesis. However, at later stages of spermatogenesis SUN4 localizes to the posterior pole of the nucleus whereas SUN3 localizes laterally along the nucleus (22, 26, 31). Moreover, in ejaculated human sperm SUN4 localizes to the midpiece (32). Sperm from SUN3- and SUN4-knockout mice show loss of attachment between the nucleus and the manchette, resulting in severe malformation of the head and consequently infertility (24, 26, 31, 33).

In contrast to SUN3 and SUN4, SUN5 localizes to the posterior part of the sperm nucleus in human and mouse during late spermatogenesis and retains this localization in ejaculated human, bovine and sheep sperm (34, 35). Both Nesprin2 and Nesprin3 were shown to be able to interact with SUN5 and the SUN5/Nesprin3 complex was shown to be important for correct localization of the proximal centriole (35–37). Furthermore, knock-out of SUN5 in mice resulted in altered Nesprin3 localization as well as loss of the sperm head-tail connection leading to infertility (25, 34–36, 38). Importantly, mutations of the SUN5 gene lead to male infertility in humans as the sperm head and tail are separated, a condition referred to as acephalic spermatozoa syndrome (ASS) (39–44, 37). ASS, despite being a rare condition, is one of the most severe forms of abnormal sperm morphology leading to infertility and affected individuals are required to undergo fertility treatment (45–48).

Here we used super-resolution microscopy to show that in mammalian sperm SUN5 is the major LINC component of the NE at the head-tail junction. Cryo-electron tomography (cryo-ET) of sperm cells thinned with cryo-focused ion beam (cryo-FIB) milling revealed a remarkably uniform spacing of 14 nm between the INM and ONM, at the base of the nucleus. We further found that this uniform membrane spacing is mediated by a regular protein lattice in the PNS. Finally, structural analysis indicated that the lattice is formed by trimers of SUN5.

## Results

### SUN5 is the major LINC component at the base of human sperm nucleus

In mammalian sperm, the head and the tail are joined at an elaborate structure called the connecting piece, which consists of several cellular elements that ensure stable coupling while still maintaining a degree of flexibility (49). We used stimulated emission depletion (STED) microscopy to determine localization of the testis-specific LINC components SUN4 and SUN5 in the connecting piece of human, boar and mouse spermatozoa. We find that SUN5 fluorescence is clearly detected at the base of the sperm nucleus in all three species **(Fig. 1A)**. Counter staining with α-tubulin and DAPI allowed to precisely define the position of SUN5 to be between the proximal centriole and the nucleus. In contrast, SUN4 primarily localizes to the midpiece of human and boar sperm **(Fig. 1B)**. This is surprising, as SUN proteins are typically localized to the NE, but is consistent with previous observations (32). In the context of the highly specialized sperm cell, SUN4 may form a non-canonical LINC complex, potentially involved in mitochondria positioning and stabilization of the midpiece. Interestingly, in mouse sperm, SUN4 signal localizes to the connecting piece around the proximal centriole like SUN5 **(Fig. 1B)**. The apparent differential localization of SUN4 in mouse compared to human and boar could be due to species-specific differences or may result from comparing different maturation stages as mouse sperm were collected from the cauda epididymis while human and boar sperm were collected from ejaculates. Overall, our fluorescence-based analysis strongly suggests that SUN5 is the main LINC complex component at the base of the nucleus in human and boar sperm.

**Fig. 1.**
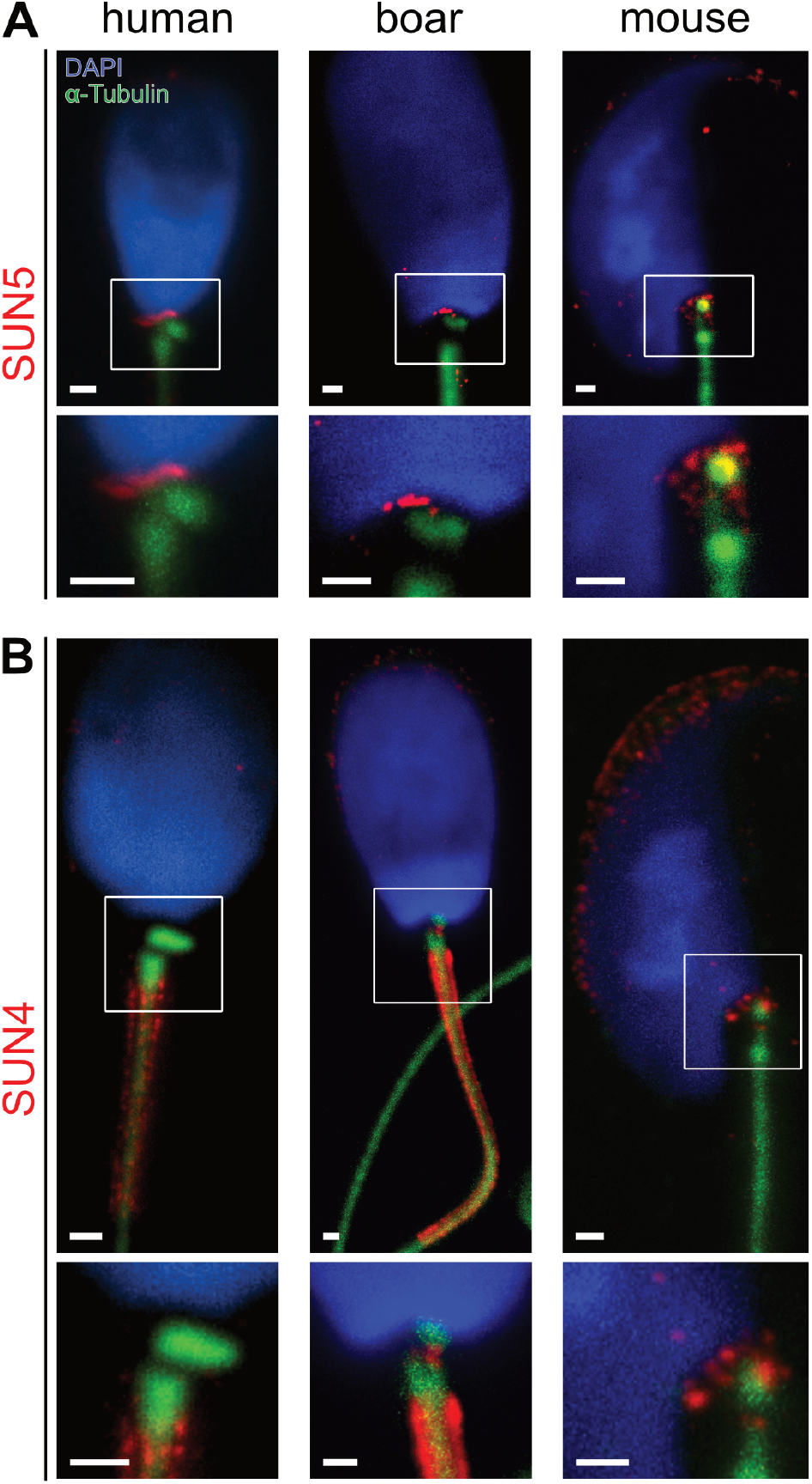
SUN5 is the major LINC component at the base of the human and boar sperm nucleus. **(A)** Stimulated emission depletion (STED) fluorescence microscopy of sperm from human, boar and mouse showing that SUN5 (red) is localized to the base of the nucleus in mammalian sperm. Counter stains, anti α-tubulin (green), and the nucleus with DAPI (blue). **(B)** STED microscopy showing that SUN4 (red) is localized to the midpiece in human and boar sperm but to the base of the nucleus in mouse. Lower panels are magnified views of the area marked with white rectangles in the upper panels. Scale bars, 500 nm.

### A conserved protein lattice maintains uniform NE spacing

To determine how SUN5-based LINC complexes are organized in the NE of human sperm, we used cellular cryo-ET. Imaging the sperm connecting piece with cryo-ET requires thinning by cryo-FIB milling, as this part of the cell is ∼800 nm thick in human sperm (for details see Material and Methods and **Fig. S1A-C**). Tomograms of the base of the head reveal common features of the sperm connecting piece, including the densely packed nuclear chromatin, the basal plate tightly aligning with the cytoplasmic side of the NE, as well as the segmented columns and parts of the proximal centriole **(Fig. 2A and B)**. The basal plate, and segmented columns consist of electron dense material without distinct structural features, similar to previous observations (50, 51). Intriguingly, we observed prominent repetitive protein densities in the NE along the basal plate **(Fig. 2B)**. Orthogonal views of the ONM reveal that the regular densities form a striking two-dimensional lattice at the perinuclear space **(Fig. 2B, Fig. S1**). At the INM, a similar protein lattice is present, albeit less pronounced **(Fig. S1D and F)**. A regular protein array at the NE of sperm from guinea pig and rat was previously described in freeze-fracture studies (52). Remarkably, the protein lattice we observe here is limited to the region at which the NE integrates with the sperm connecting piece, like the SUN5 fluorescence signal. At sites where the NE bends towards the postacrosomal sheath we instead observe clusters of nuclear pore complexes, consistent with previous findings (53) **(Fig. 2C)**. Furthermore, the lattice is not observed in the NE near the postacrosomal sheath and acrosome **(Fig. 2D and E)**. Within the postacrosomal sheath regularly arranged membrane protein densities occurs outside the NE as was previously described for boar sperm (54).

**Fig. 2.**
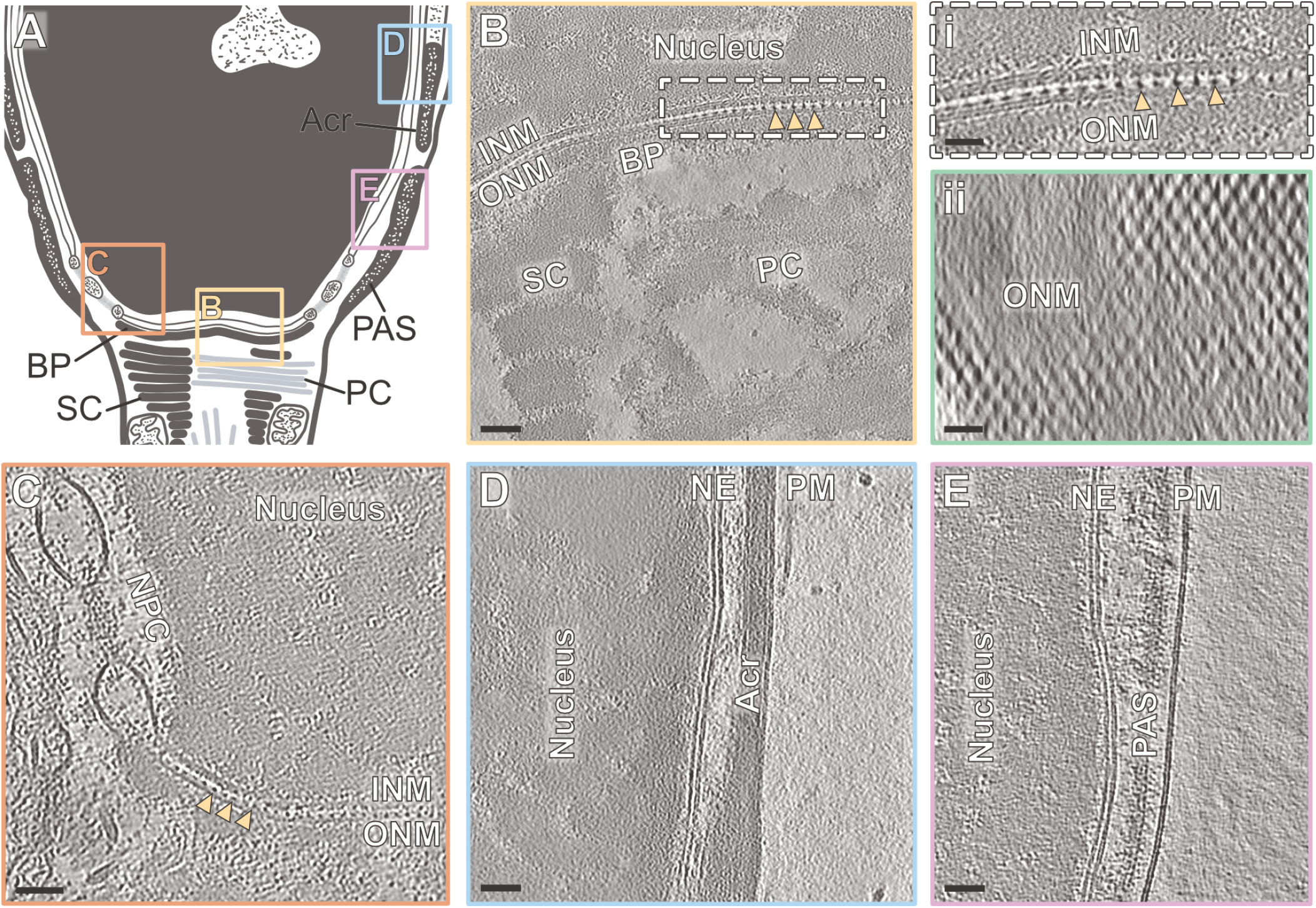
A LINC lattice limited to the base of the nucleus of human sperm. **(A)** Schematic depiction of human sperm head and connecting piece. Structures in the connecting piece are annotated, including the basal plate (BP), segmented columns (SC), proximal centriole (PC), acrosome (Acr) and postacrosomal sheath (PAS). Colored rectangles highlight the areas imaged with cryo-ET. **(B)** A tomographic slice of the connecting piece. A regular protein lattice (yellow triangles) between the INM and ONM coincides with the basal plate (BP). The segmented columns (SC) and proximal centriole (PC) are also visible. The marked area is magnified in **(i)** and an orthogonal view (4.3 nm thick slice) of the ONM **(ii)** highlights the 2D arrangement of the lattice. **(C)** A tomographic slice capturing the base and side of the nucleus, revealing a regular protein lattice (yellow triangles) between the INM and the ONM ending just before a nuclear pore complex (NPC). **(D)** A tomographic slice close to the equatorial region capturing the nuclear envelop (NE), acrosome (Acr) and plasma membrane (PM). **(E)** A tomographic slice of the postacrosomal sheath (PAS). Scale bars, (B-E) 50 nm, (i, ii) 25 nm.

Inspecting multiple tomograms capturing the NE along the base of the sperm nucleus indicated that the spacing between INM and ONM is strikingly uniform and tight (13.75 +/-1.29 nm) when the protein lattice is present **(Fig. 3A and F)**. Of note, the membrane spacing observed here is significantly shorter than the membrane spacing reported for other cell types, for example C. elegans (∼50 nm), Xenopus oocytes (∼30 nm) and yeast (∼25 nm) (55–57). Interestingly, a recent study reported even tighter membrane spacing in the human sperm NE (∼9 nm) for areas distal from nuclear pore complexes and wider spacing (∼27 nm) in areas closer to NPCs (53). In contrast to the membrane spacing observed at the base of the head, we find the spacing to be ∼5.5 nm near the postacrosomal sheath and acrosome **(Fig. 2D and E)**. This indicates that the LINC lattice is involved in maintaining a uniform membrane spacing of ∼14 nm along the human sperm connecting piece.

**Fig. 3.**
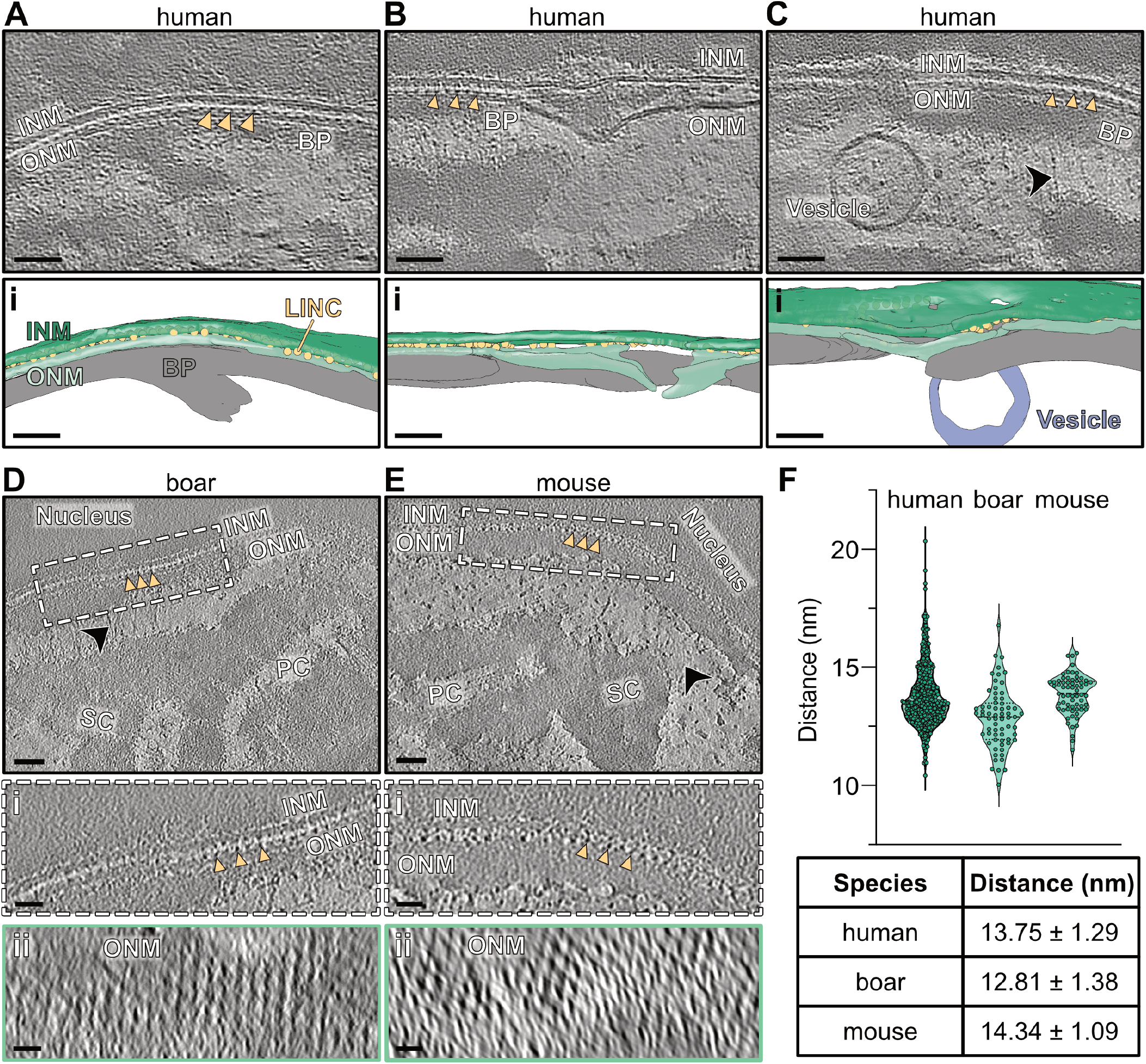
The LINC lattice is associated with a uniform membrane spacing and is conserved in mammals. (A-C) Tomographic slices of the connecting piece of different human sperm cells, with 3D segmentations below **(i)**. Notably the membrane spacing is uniform when the lattice is present (yellow triangles). Filaments (black arrowheads) extending from the basal plate (BP) are reaching the upper part of the segmented columns. **(D-E)** Tomographic slices of the boar **(D)** and mouse **(E)** sperm connecting piece, displaying a similar LINC lattice in the NE as observed in human sperm (yellow triangles). The marked areas are magnified in **(i)** and an orthogonal view (4.3 nm thick slice) of the ONM is shown in **(ii)** of the corresponding tomographic slice. **(F)** Quantification of the distance between the ONM and the INM for the different species. The table shows mean distance and standard deviation of the measurements. Scale bars, (A-E) 50 nm, (i, ii) 25nm.

LINC complexes have previously been hypothesized to serve as “molecular rulers” and to stabilize uniform membrane spacing of the NE depending on the lengths of their coiled-coil domains (10, 20, 58). Functionally, the regulation of membrane spacing by LINC complexes is crucial for ensuring nuclear morphology in cells exposed to physical stress, as has been shown in force-bearing muscle cells in C. elegans (55). As the connecting piece of sperm cells must withstand forces exerted by flagellar beating, the protein array observed at the NE may play a role in maintaining nuclear morphology and attachment to the tail.

Interestingly, we also observed areas with irregular membrane spacing and bulging of the ONM in seven out of twelve tomograms and vesicles were located on the cytoplasmic side near the bulging ONM in three tomograms **(Fig. 3B and C, Fig. S2)**. Consistent with the observation that the uniform membrane spacing coincides with the protein lattice, its presence is drastically reduced at sites of membrane bulging. Moreover, the basal plate was interrupted at sites where membrane bulging occurred, indicating that the lattice may also be tethering the basal plate to the NE. On the cytoplasmic side, we find filaments extending from the basal plate to the upper part of the segmented columns, sometimes referred to as the capitulum **(Fig. 3C)**, which suggests a molecular network coupling the segmented columns, basal plate and the nucleus.

As our fluorescence data indicated that SUN5 is also present in the connecting piece of boar and mouse sperm, we next imaged the nucleus along the connecting piece of sperm from these species. Like our in situ data from human sperm, the tomograms show a regular protein lattice along the NE that appears to maintain uniform membrane spacing **(Fig. 3D-F)**. Moreover, filaments similar as those observed in human sperm connecting the basal plate to the upper part of the segmented columns are present in both boar and mouse sperm. This implies that despite the significant differences in head morphology between these three species, the molecular mechanism of head-tail attachment including the LINC lattice in the NE is conserved.

### In situ 3D reconstruction of the LINC lattice

To analyze the organization of the putative SUN5 lattice, we used subtomogram averaging. An initial reconstruction from one tomogram showed a clear pattern along the ONM with six densities evenly distributed around a central density and consistently spaced 12.4 nm apart **(Fig. S3A)**. Likewise, regularly spaced densities were also apparent at the INM directly opposing the ONM densities. Using this initial reconstruction as a reference, more particles were picked by template matching and subsequently combined, resulting in a similar lattice structure resolved to ∼27 Å **(Fig. 4A and B, for details see Material and Methods)**. In addition to the main lattice, this reconstruction revealed a very faint density orthogonally connecting the opposing ONM and INM densities **(Fig. 4A)**. Moreover, each density in the lattice along the ONM appears to form lateral connections to the neighboring density **(Fig. 4A)**. Similar structural analyses for boar and mouse sperm resulted in comparable lattice structures along the ONM and INM with a spacing of 12.3 nm and 12.6 nm, respectively **(Fig. S3B and C)**. The 3D reconstructions of the lattice further show a spacing between ONM and INM of 13.5 nm, consistent with the direct measurements from the tomograms **(Fig. 3F and 4A, Fig. S3)**.

**Fig. 4.**
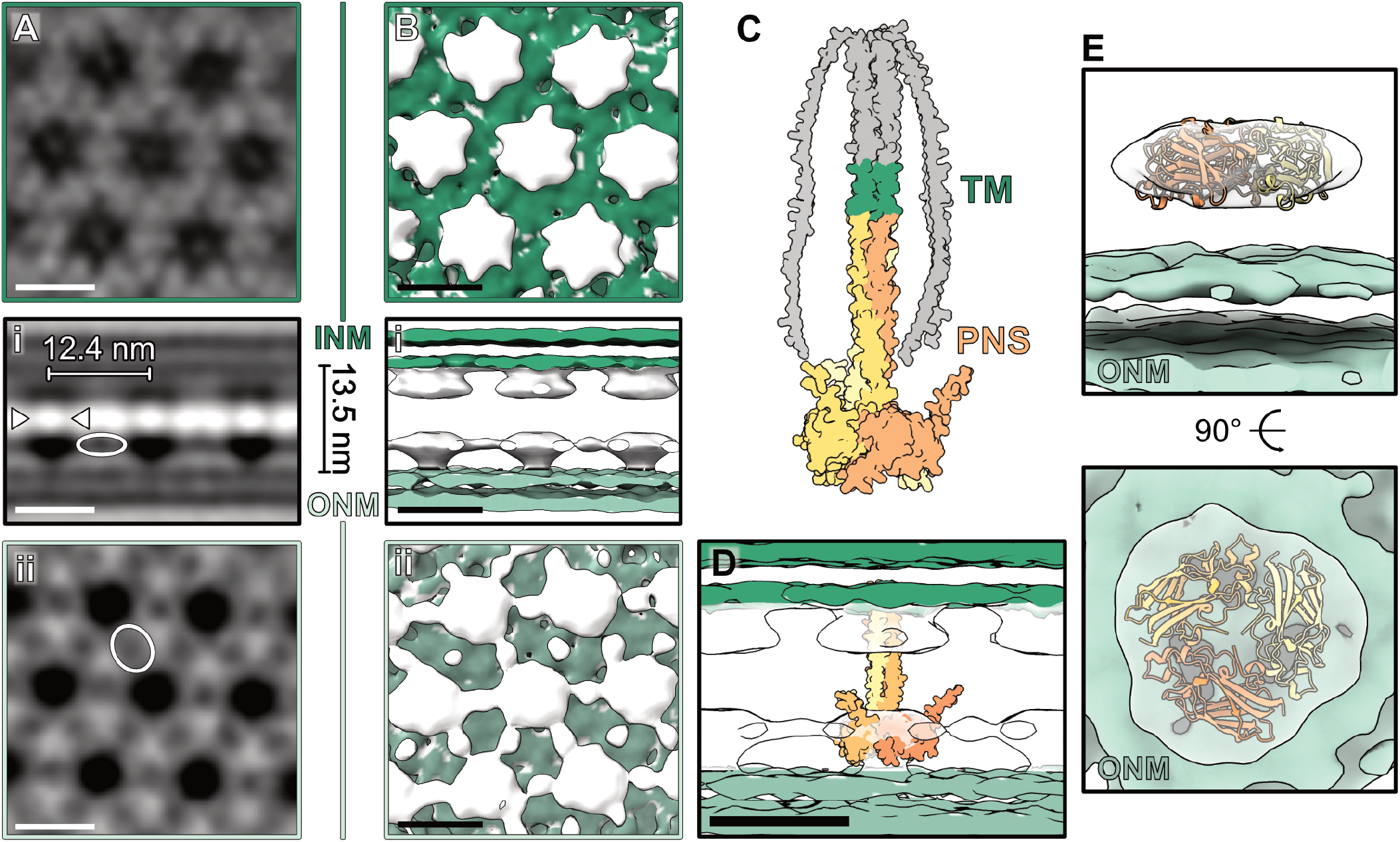
In situ 3D reconstruction of the LINC lattice. **(A)** 2 nm thick slices through the 3D reconstruction (lowpass-filtered to 27 Å) of the LINC lattice along the INM **(A)**, the NE membranes **(i)** and the ONM **(ii)**. Note the faint connections between the INM and ONM densities (white arrows) and lateral connections between ONM densities (white circles). **(B)** Isosurface representation of the reconstruction of the LINC lattice with views corresponding to the panels in (A). **(C)** AlphaFold3 prediction of a full-length human SUN5 trimer. The predicted peri-nuclear space portion (PNS) are colored in different shades of yellow and the transmembrane helices (TM) are in green. **(D)** Fit of the predicted peri-nuclear space portion of human SUN5 (123-379, yellow) into the 3D reconstruction of the LINC lattice shown in **(B). (E)** Isosurface representation of a reconstruction of one central density at the ONM with C3 symmetry applied and a fit of a trimeric SUN5 domain (205-364, yellow). Different threshold levels were used to visualize the ONM bilayer (3.11) and the protein density (7.08). Scale bars, 10 nm.

Current models of LINC complex topology suggest that the transmembrane part of the SUN proteins is embedded in the INM, followed by a coiled-coil region stretching across the perinuclear space to allow SUN domain interaction with the C-terminal part of a Nesprin protein at the ONM (8, 11–14, 20). According to this model, the density observed at the ONM would represent SUN domains. In vitro, different oligomeric assemblies of SUN domains from SUN1 and SUN2 proteins in complex with a Nesprin C-terminus peptide were reported, including trimers, dimers of trimers and trimers of trimers (17–19). This prompted the question which oligomeric state is formed in situ in the context of the lattice in the sperm NE. To answer that question, we attempted fitting available crystal structures of the different SUN domain assemblies into our reconstruction from human sperm. Only a single SUN domain trimer produced a reasonable fit for the ONM density **(Fig. S4A)**. In contrast, dimeric or trimeric assemblies of SUN domain trimers are significantly larger than the resolved density **(Fig. S4B and C)**. Consistent with this, an AlphaFold prediction of a trimer of the PNS-localized portion of SUN5, including the coiled-coil region and SUN domain (residues 123-379), concurs with the observed uniform membrane spacing between ONM and INM of ∼13.5 nm **(Fig. 4C and D)**. While AlphaFold predictions of SUN3 and SUN4 produced similar fits, predictions of SUN1 and SUN2 display a significantly longer coiled-coil region **(Fig. S5C)**, consistent with previous observations suggesting that SUN1 and SUN2 maintain membrane spacing at 40-50 nm (20, 58). Based on our conclusion that the ONM densities represent SUN5 SUN domain trimers, we next generated a focused reconstruction of a single ONM density with C3 symmetry applied. This resulted in a 20.1 Å resolution reconstruction to which we could fit an AlphaFold predicted structure of a SUN5 SUN domain trimer **(Fig. 4E, Fig. S3E)**.

## Discussion

Combining existing knowledge on LINC complex composition with our fluorescence data, in situ 3D reconstruction, and structure predictions, we propose a model in which SUN5 trimers assemble into a 2D lattice along the NE at the base of the sperm head **(Fig. 5)**. In this model SUN5 forms a trimer in the INM, with its N-terminus interacting with the nuclear lamina via Lamin B1 (38, 59). Within the PNS, SUN5 stretches its coiled-coil region so the SUN domain can interact with the C-term of a Nesprin protein that is embedded in the ONM. The major portion of a Nesprin protein **(Fig. S6A and B)** localizes to the cytoplasm and is likely part of the basal plate. Previous studies reported direct interactions between SUN5 and Nesprin3 in human and mouse sperm (36, 37). Another study described the formation of a Nesprin2/SUN5 complex in mouse and human during the meiosis stages of spermatogenesis (35). As our study focused on ejaculated sperm, Nesprin3 appears as the most probable candidate to interact with SUN5. Finally, filaments connect the basal plate to the upper portion of the segmented columns, completing an extensive interaction network reaching from the nuclear lamina, across the NE (via LINC complexes) into the connecting piece **(Fig. 5)**.

**Fig. 5.**
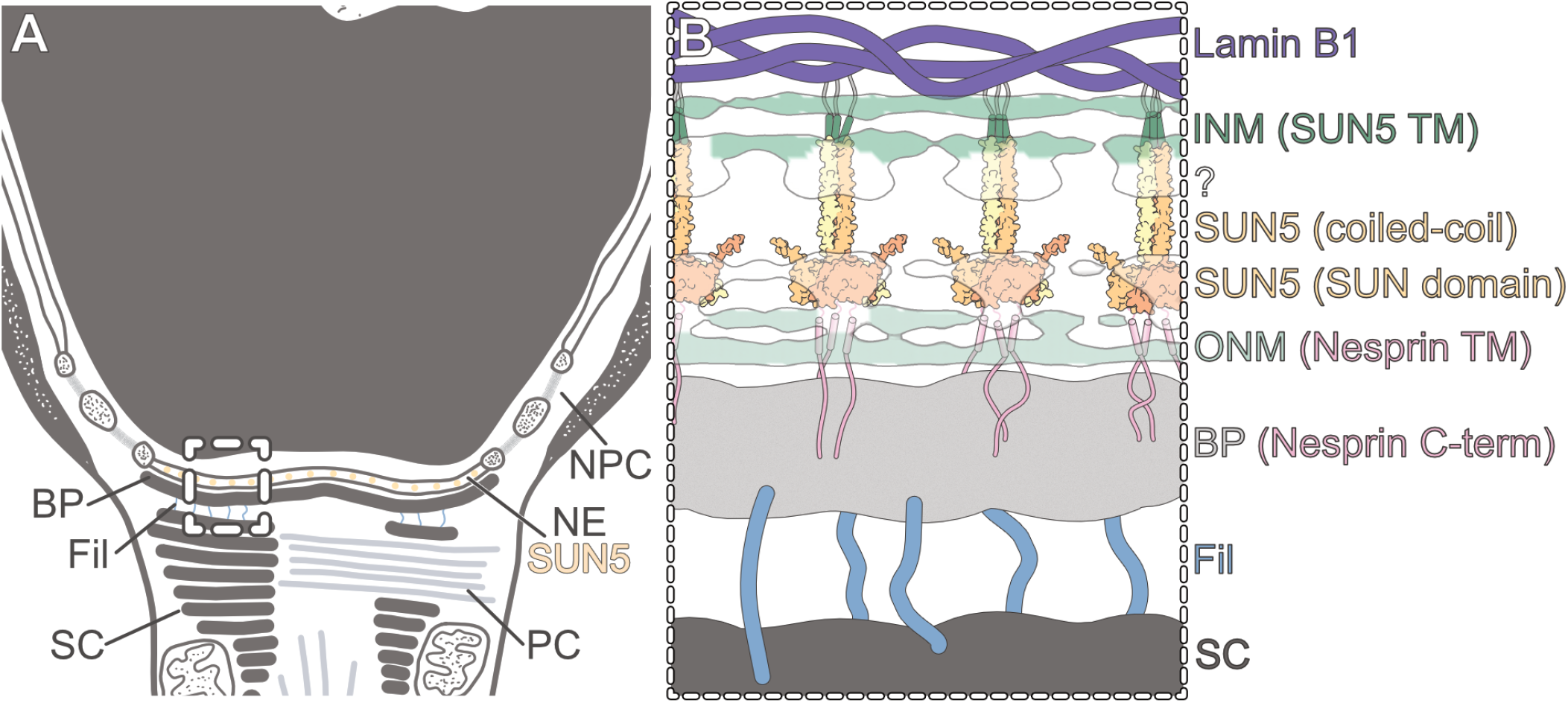
Proposed model for the molecular network at the base of the sperm head. **(A)** Schematic of the sperm connecting piece highlighting SUN5 LINC complexes in the NE along the basal plate (BP) and filaments (Fil) connecting the BP and upper part of the segmented columns (SC). **(B)** Zoom-in on the area marked in (A). The organization of SUN5 (yellow) between the ONM and INM establishes a bridge between nuclear lamina (Lamin B1) and BP, which is directly connected to the segmented columns via filaments.

Our data clearly show that the LINC complexes assemble into a lattice but the driving forces for this higher-order assembly remain elusive. Intriguingly, structure prediction of full-length SUN5 suggests C-terminal extensions that may be involved in lateral interactions between the SUN domain trimers **(Fig. 4C and D)**. These extensions are unstructured and ∼7 nm long, allowing them to bridge approximately half the distance of the average lattice spacing (∼12.4 nm) and can potentially account for the densities we resolved between SUN domain trimers **(Fig. 4A)**. Alternatively, directly interacting Nesprin proteins could play a role in the lattice assembly by connecting individual SUN trimers as suggested before (8). It is further possible that other interacting proteins are needed for the lattice assembly. For instance, cytoskeletal actin was reported to assemble SUN2/Nesprin2 LINC complexes into linear arrays (60). Moreover, multiple studies have identified SUN5 interacting proteins, including CENTLEIN and Septin12. CENTLEIN is a protein required for centriole cohesion in somatic cells and was shown to directly interacts with SUN5 during spermatogenesis in mouse to bridge the interaction with PMFBP1 (61, 62). Septin12, a cytoskeletal GTPase that may restrict membrane protein diffusion at the NE, directly interacts with SUN5 in immunoprecipitation experiments and co-localizes with SUN5 in epididymal mouse sperm (38, 53).

In our lattice reconstruction, we observed a repetitive unassigned density at the INM following the same lattice parameters as the density at the ONM. This density could be part of SUN5 itself, provided the part proximal to the INM adapts a different conformation than the predicted coiledcoil. Alternatively, other proteins could be interacting with the SUN5 coiled-coil region close to the INM. For instance, the INM protein Emerin interacts with LINC complexes in somatic cells, although no direct interaction with SUN5 has been reported (5, 6, 28). Ultimately, identifying the precise composition of the full LINC lattice, particularly the lateral connections between SUN domain trimers as well as the density observed at the INM, will require further studies.

In summary, by combining super-resolution fluorescence microscopy and in situ cryo-ET, we find that SUN5 localizes to the base of the mammalian sperm nucleus, where it is involved in forming an extensive hexagonal lattice. Interestingly, the lattice is absent in areas where ONM bulging is observed and the BP is interrupted, suggesting a role of the lattice in tethering the two nuclear membranes to each other like molecular velcro and thereby maintaining stable and uniform membrane spacing. The potential interaction network between nuclear lamina, nuclear envelope and basal plate – mediated by the LINC lattice – would also provide a structural basis for how SUN5 may be involved in anchoring the sperm head to the tail, rationalizing how mutations in SUN5 result in ASS (45–48). Considering the role of canonical LINC complexes in mechanotransduction, the LINC lattice may additionally be involved in transmitting signals from the cytoplasm to the nucleus. For instance, changes in motility induced during capacitation could trigger chromatin remodeling observed after capacitation of human sperm (63). Whether the LINC lattice indeed has roles beyond structurally reinforcing the head-tail junction remains to be elucidated.

## Acknowledgements

We thank Brigitte Arends for organizing and providing donor sperm samples. T.Z. was funded by the European Research Council (ERC-2022-COG project 101088673) and J. M. by the Marie Skłodowska-Curie grant (No 101153233).

## Author Contributions

TZBM, JM, SD and MRL designed research. JM, SD and MRL performed research and analyzed data. TZBM and JM wrote the paper and all authors contributed to the final version of the paper.

## Materials and Methods

### Sperm preparation

Ejaculated human donor sperm was either received frozen in straws from the Utrecht University Medical Center (protocol number 21-754) and stored in liquid nitrogen or nitrogen vapor. Immediately before the start of an experiment, straws were thawed for 30 s in a 37°C water bath. Subsequently, the sperm sample was layered onto a discontinuous two-step Percoll gradient (35 %, 70 %). Percoll dilutions were prepared in HBS buffer (20 mM HEPES (pH 7.6), 137 mM NaCl, 10 mM glucose, 2.5 mM KCl). The gradient was centrifuged for 10 min at 300 g followed by another centrifugation step for 20 min at 750 g to isolate motile sperm. The pelleted motile sperm was resuspended in DPBS, the cell concentration determined in a counting chamber and the cells diluted to the desired concentration. Boar sperm was obtained commercially (Varkens KI Nederland) and received and stored at 18°C in an extender solution. Processing of the sample was carried out within 24 h after delivery of the sample. Boar sperm was applied to a Percoll gradient and further processed as described for human sperm. Mouse (Mus musculus) sperm was isolated from the cauda region of the epididymis of C57BL/6 mice as previously described (64). Caudal mouse sperm was kept in a modified Biggers, Whitten, and Whittingham media (BWW: 20 mM HEPES, 91.5 mM NaCl, 4.6 mM KCl, 1.7 mM D-glucose, 0.27 mM sodium pyruvate, 44 mM sodium lactate, 5 U/mL penicillin, and 5 µg/mL streptomycin, adjusted to pH 7.4 and an osmolarity of 300 mOsm/kg) and was washed by centrifugation at 400 g for 15 min through a 27 % Percoll layer followed by another centrifugation (400 g, 2 min) to remove excess Percoll.

### Cryo-EM grid preparation

Human sperm samples were diluted to 10-20 × 10^6^ cells/mL in DPBS or DPBS containing 5 % glycerol. Mouse and boar sperm samples were diluted to 20-30 × 10^6^ cells/mL. 3.0-3.5 µL of the diluted cell suspension were loaded onto a glow-discharged Quantifoil R2/1 or R2/2 200 mesh holey carbon grid. The grids were blotted from the back for 5-6 s before plunging into liquid ethane cooled to liquid nitrogen temperature. Plunge-freezing was carried out using a manual plunger (MPI Martinsried) or an EM GP2 Automatic Plunge Freezer (Leica). Afterwards, grids were clipped into Autogrids (Thermo Fisher Scientific) for mechanical support and stored in liquid nitrogen.

### Cryo-FIB milling

Grids were loaded into an Aquilos (Thermo Fisher Scientific) cryo-focused ion beam/scanning electron microscope. Grids were mapped and monitored during milling using the SEM at 2 kV and 13 pA. Grid maps were collected before and after deposition of platinum GIS for 6-8 s. Subsequently, grids were sputter coated with platinum (1 kV, 30 mA, 10 Pa, and 10 seconds). Targeting with the FIB was performed at 30 kV and 1.5 or 10 pA. Samples were milled at an effective milling angle of 11-12° in four consecutive steps: a rough milling step at 0.5 or 1 nA, including milling of micro expansions for stress-relief (65), an intermediate milling step at 0.1 or 0.3 nA, a fine milling step at 50 pA or 0.1 nA and a polishing step at 30 pA. Polished lamellae were stored in liquid nitrogen until imaging.

### Cryo-ET data collection

All data was collected on a 200 kV Talos Arctica (Thermo Fisher Scientific) microscope equipped with a post-column energy filter (Gatan) and a K2 Summit direct electron detector (Gatan). Data collection was carried out in zero-loss imaging mode with a 20 eV slit width using SerialEM (66). For lamellae from human sperm, images were acquired following a grouped dose-symmetric tilt scheme covering a range from -48° to 48° or -56° to 56° in 4° increments and a maximum total dose of ∼100 e-/Å^2^. Initial overview tilt series were collected at a pixel size of 3.57 Å and a target defocus of -6 µm and tilt series for subtomogram averaging were collected at a pixel size of 2.17 Å and a target defocus of -3 to -5 µm. Mouse and boar sperm data were collected similarly but with an effective tilt range of -56° to 56° or -54° to 54° with a 2° or 3° tilt increment, respectively. The defocus was set between -4 to -6 µm and frames were recorded at a pixel size of 2.17 Å (boar sperm) or 2.83 Å (mouse sperm).

### Tomogram reconstruction

Individual frames were aligned either on-the-fly using WARP or with MotionCor2/1.6.4 (67, 68). For segmentation and visualization, four times binned tomograms were reconstructed with AreTomo1.3.4 and subsequently denoised with cryo-CARE 0.3.0 (69, 70). For subtomogram averaging, tomograms were reconstructed using IMOD 4.12.62 by weighted back-projection and tomograms were CTF-corrected using *ctfphaseflip* (71).

### Subtomogram averaging

Subtomogram averaging was carried out in PEET (1.18.0) with missing wedge compensation and based on IMOD-reconstructed tomograms (72, 73). A subset of positions for subtomogram extraction was initially defined by manually placing two model points per particle, one at the ONM and one at the INM (bin4, pixel size 8.68 Å). The *stalkInit* function of PEET was used to generate initial motive list and rotation axis files, defining the initial particle orientation. Subtomograms of approximately 60 nm x 40 nm x 60 nm were extracted and computationally aligned. For the alignment, a cylindrical soft-edged mask with a height of 14 nm and a radius of 20 nm was applied to mask out the nuclear membranes. A similar masking strategy was applied during all subsequent steps. Based on the known orientation of the particles relative to the two membranes, the initial alignment allowed for a full search around the y axis (defined to be perpendicular to the membranes) and 30° search ranges around the x and z axis. The initial alignment resulted in an average volume displaying six densities oriented around a central density at the ONM. Similar results were obtained from different tomograms, and a resulting averaged volume was used as a reference in following alignments. To define particle positions in all remaining tomograms, the ONM was segmented manually in IMOD and points were seeded along the meshed surface using *meshInit* at half the distance at which densities at the ONM were observed in the initial averages. Particle positions derived from seeding with *meshInit* were then refined by template matching to the reference volume from the initial alignment. If necessary, patches of NE not covered by this strategy were filled by using *modifyMotiveList* to create particles with a shifted position compared to the existing ones. Particle shifts were based on the observed distances between densities at the ONM. The resulting particle positions were extracted using *createAlignedModel* and another alignment was carried out. Duplicate particles as well as particles with low cross-correlation were removed using PEET extension scripts (kindly provided by Dr. Daven Vasishtan) before combining data from different tomograms. This resulted in a total of 4501 particles from eight different tomograms, each tomogram capturing a different sperm cell. Particle positions were randomized around their y-axis to reduce bias potentially introduced during template matching. Based on the initial observation of 6 densities surrounding a central density, the search ranges for refinement were set to 64° around the y axis and 30° around the x and z axis. Particle heterogeneity was assessed using principal component analysis followed by k-means clustering (*pca* and *clusterPca* functions in PEET), and particles not showing significant density at the INM and ONM were removed, resulting in the final number of 3788 particles. The refinement was repeated with these particles, and the resulting averaged volume was used as a new reference for further refinement steps. Finally, particles were re-extracted from two times binned tomograms (pixel size 4.34 Å) and another alignment using the same search ranges as above was carried out. To focus on a single nuclear envelope density, particle coordinates from the final refinement described above were used with an adjusted box size of approximately 14 nm x 22 nm x 14 nm and the radius of the mask was reduced to 5 nm. Particles were reextracted from unbinned tomograms and C3 symmetry was applied by using a PEET extension script automatically rotating the original particles by 120° around the y axis. A new reference of the symmetrized particles was created by averaging all original and virtual particles before running another alignment to create the final averaged volume. For all averaged volumes, FSC curves were determined by splitting the dataset into even and odd particles, running separate alignments and using *simpleFSC* with the same mask applied as during the alignment. Details on cryo-ET data collection and subtomogram averaging are summarized in Table S1.

### Segmentation and visualization of tomograms

Generally, cryo-CARE denoised tomograms were used for segmentation and visualization. The nuclear envelope was segmented using Membrain v2 and the resulting segmentation manually cleaned in UCSF ChimeraX (74, 75). The basal plate was segmented manually using the drawing tools in IMOD and the resulting model was smoothened using *mtffilter*. To segment the nuclear envelope densities, the model files produced for particle extraction in subtomogram averaging after refinement were loaded in the ArtiaX plugin in ChimeraX and displayed as spheres (76). Segmentations were combined and visualized in ChimeraX. Representative tomogram slices were captured with IMOD.

### Distance measurements and quantification

Distances were measured in IMOD by placing two model points at the distance supposed to be measured. At least 50 measurement points per tomogram were placed to derive the intermembrane spacing directly from eight different tomograms. Intermembrane spacing from subtomogram averages were measured similarly, with a minimum of 8 measurements along the membrane. Points were placed directly on the PNS-facing lipid layer and only at positions where both the INM and ONM bilayer of the nuclear envelope were clearly discernible. Spacing of individual densities within the lattice was derived by placing six pairs of points spanning from the central particle to the surrounding particles for both the INM and ONM densities. Distances were obtained by running *imodinfo* on the corresponding model files and plotted using GraphPad Prism.

### Transmembrane region and AlphaFold prediction

For SUN5, three copies of the full-length protein were submitted for structure prediction by AlphaFold3, resulting in the successful prediction of a trimer (77). Subsequently, DeepTMHMM was used to predict the transmembrane helices of all human SUN proteins (78). Based on the determined transmembrane helices AlphaFold3 was used to predict the structure of trimers of only the PNS-localized portion of all five human SUN proteins **(Fig. S5**). The resulting models were fitted in ChimeraX, orienting the SUN domains based on the density at the ONM. For the prediction of a SUN5 SUN domain trimer, only residues 205-364 were submitted for prediction. All models were visualized and assessed in ChimeraX.

### Immunofluorescence staining and microscopy

For immunofluorescence (IF) staining of sperm cells, 1.5 coverslips (CARL ROTH, 53586) were pre-coated with 200 µL of poly-L-lysine (Sigma, P4832) and incubated for 15 minutes before removing excess liquid and allowing them to dry. Sperm samples were thawed and washed, after which 100–200 µL of the cell suspension was pipetted onto the coated coverslips and incubated for 30-60 min to allow cell adhesion. Excess liquid was then removed, and fixation was performed by adding 200 µL of 4 % formaldehyde (Thermo Fisher Scientific, FB002), incubating for 15 minutes at room temperature (RT) under a fume hood, followed by two washes with 400 µL of DPBS. Permeabilization was carried out by incubating the coverslips in 300 µL of DPBS containing 0.5 % Triton X-100 for 5 minutes at RT, followed by two additional washes with 400 µL of DPBS. Blocking was performed by incubating the coverslips in 400 µL of blocking solution (DPBS with 10 % BSA) for 1 hour at RT. Primary antibody staining was conducted by incubating the coverslips overnight at 4°C in 300 µL of blocking solution containing the following primary antibodies: SUN5 (Proteintech, 17495-1-AP) at 1:100 for human and 1:50 for mouse, primary conjugated Alexa 488 alpha Tubulin Monoclonal Antibody (B-5-1-2) (Thermo Fisher Scientific, 32-2500) at 1:250, SUN4 (Proteintech, 19721-1-AP) at 1:100 for human and 1:50 for mouse. After primary antibody incubation, the coverslips were washed three times for 5 minutes each with 400 µL of washing solution (DPBS + 0.05 % Tween-20) on a gentle shaker. Subsequently, coverslips were incubated for 2 hours at RT with gentle shaking in 300 µL of secondary antibody solution, which included anti-rabbit Alexa 594 (Thermo Fisher Scientific, A-21207) at 1:200, Atto 647N (Sigma, 50185) at 1:200, and DAPI (Thermo Fisher Scientific, 62248) at 1:1000. After incubation, coverslips were washed three times for 5 minutes each with 400 µL of washing solution to ensure thorough removal of unbound antibodies. Coverslips were then carefully lifted and allowed to dry for a few minutes before mounting. A drop of mounting medium (Thermo Fisher Scientific, P36965) was applied to a glass slide, and the dried coverslip was placed onto it, with the sample facing the mounting medium. Mounted samples were stored at 4°C for 24 hours prior to imaging. Super-resolution STED imaging was performed using the Abberior MINFLUX system with a 60x oil objective. Throughout the procedure, samples and antibodies were protected from light to preserve fluorescence signal integrity.

## Supplementary Information

**Fig. S1.**
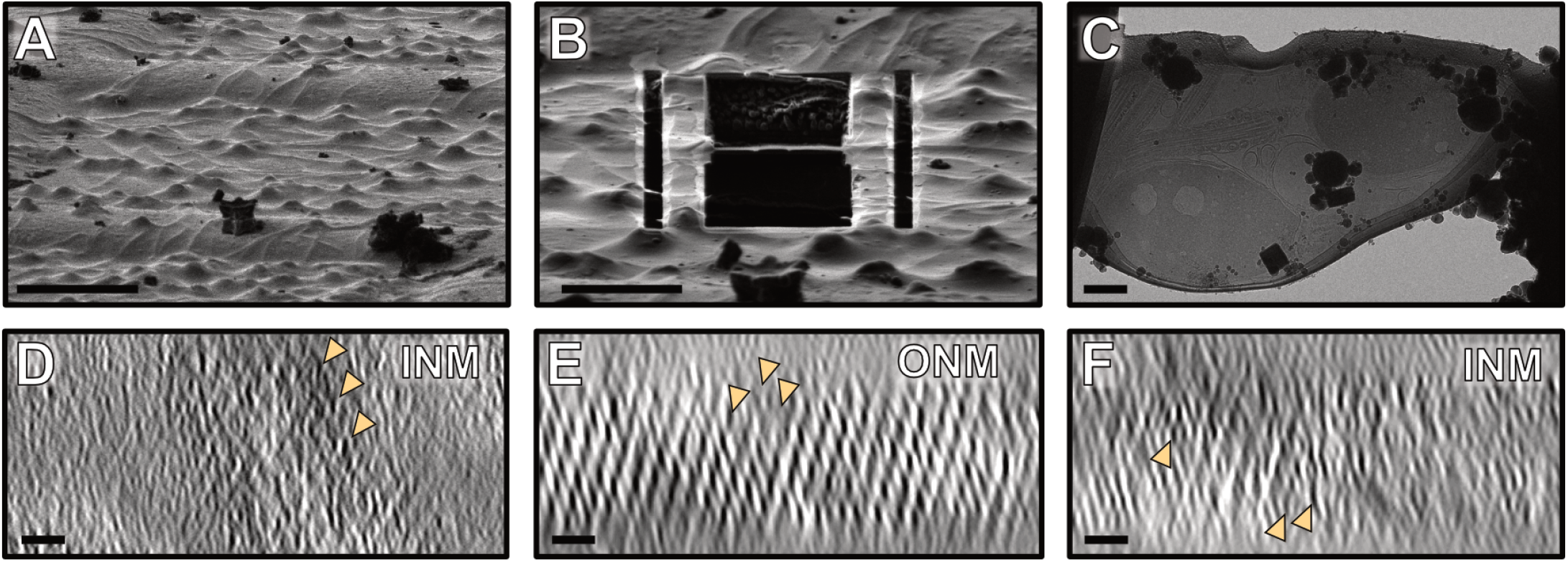
FIB milling provides access to the sperm connecting piece. **(A-B)** Ion beam snapshots of a grid with human sperm before **(A)** and after **(B)** cryo-FIB milling. **(C)** An overview image of a cryo-FIB lamellae containing two human sperm heads, the connecting piece and parts of the tails. **(D)** Orthogonal view at the INM of the tomogram shown in Fig. 2E. **(E-F)** Orthogonal views at the ONM **(E)** and INM **(F)** of the tomogram shown in Fig. 2D. Scale bars, (A) 20 µm, (B) 10 µm, (C) 1 µm, (D-F) 25 nm.

**Fig. S2.**
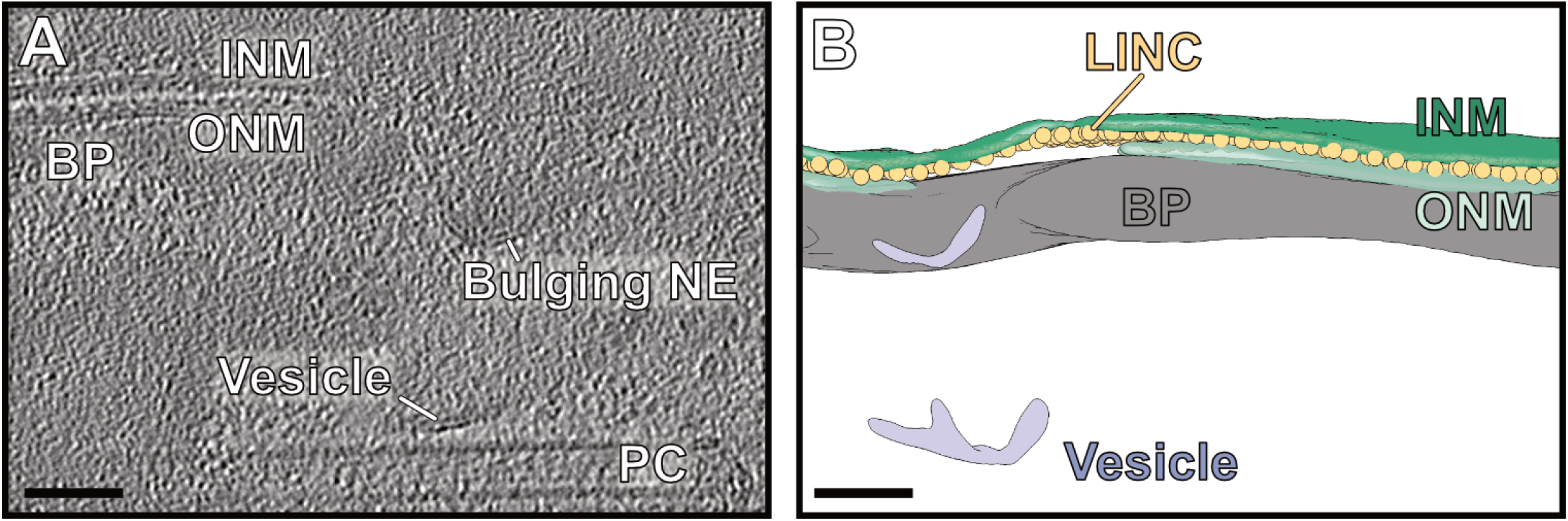
Uniform membrane spacing is only maintained in presence of the LINC lattice. **(A)** A tomographic slice of the sperm connecting piece showing ONM bulging and a vesicle near the bulging ONM. **(B)** Segmentation of the INM, ONM, BP, vesicular membranes and LINC complexes of the tomogram shown in **(A)**. Scale bars, 50 nm.

**Fig. S3.**
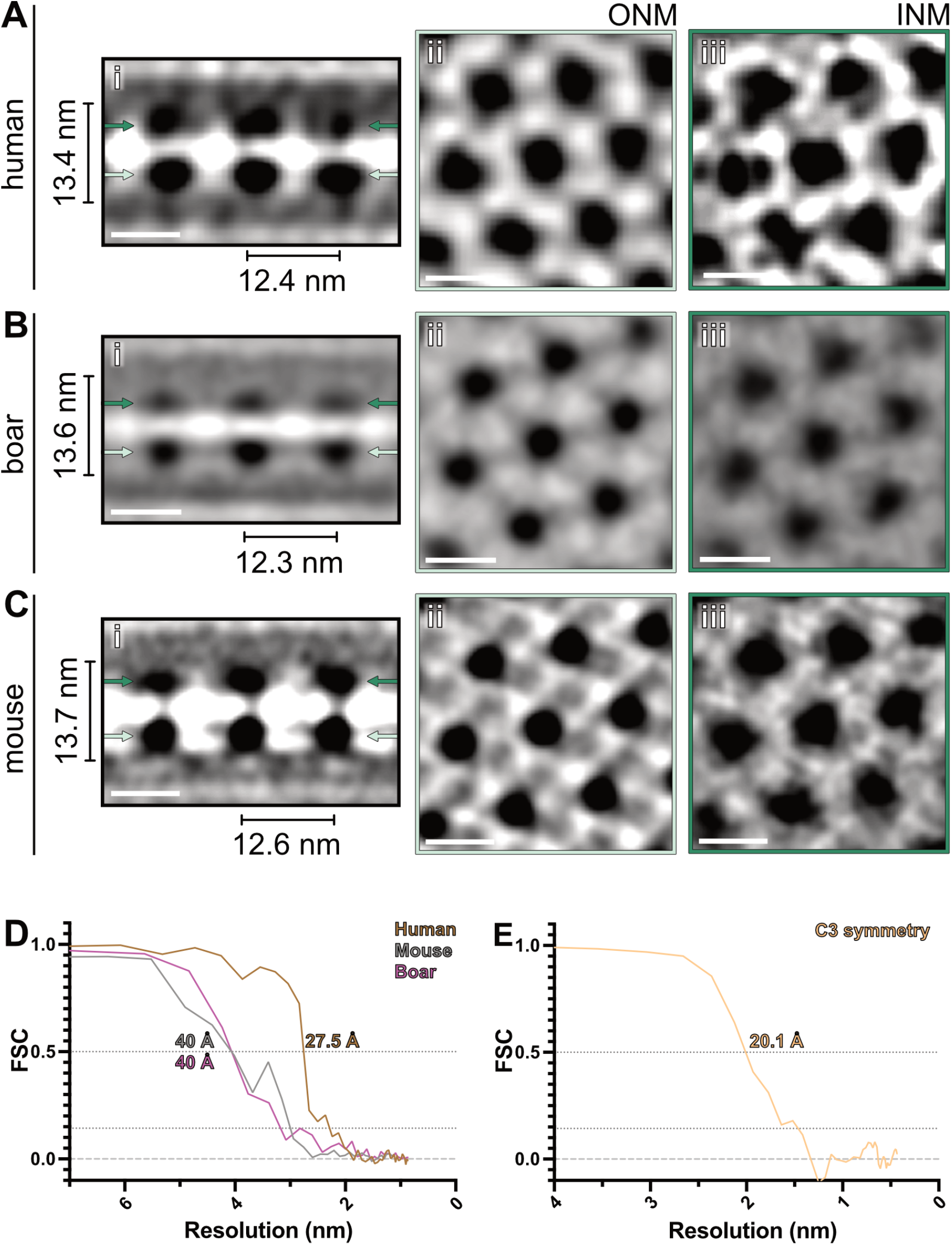
SUN5 lattice geometry is conserved in human, boar and mouse sperm. **(A-C)** Side views **(i)** and orthogonal views at the ONM **(ii)** and INM **(iii)** of the 3D reconstruction of the SUN5 lattice from human (A), boar (B) and mouse (C) sperm. The 3D reconstruction from human sperm presented in **(A)** is the initial reference produced from 133 manually picked particles. No symmetry was applied for these 3D reconstructions, and they were lowpass-filtered to 40 Å. **(D)** FSC plots for the 3D reconstruction from mouse and boar sperm and of the final 3D reconstruction from human sperm (Fig. 4A, B). **(E)** FSC plot for the 3D reconstruction of a single ONM particle with C3 symmetry applied (Fig. 4E). Scale bars, 10 nm.

**Fig. S4.**
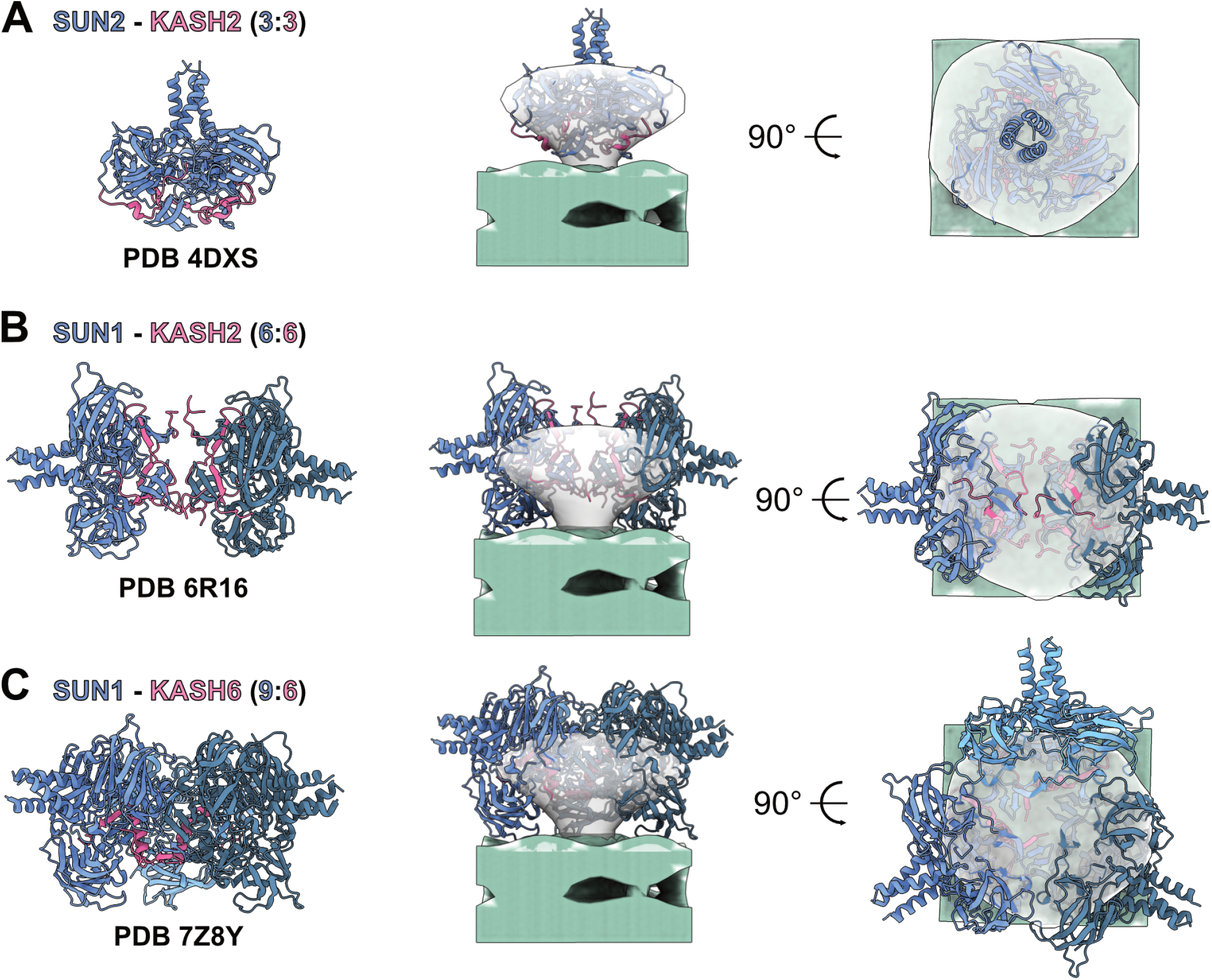
A SUN domain trimer fits the density observed at the ONM. **(A-C)** Crystal structures of a SUN2-KASH2 trimer **(A)**, a SUN1-KASH4 dimer of trimers **(B)** and a SUN1-KASH6 trimer of SUN domain trimers bound to six KASH6 peptides **(C)** and the corresponding fits into the density observed at the ONM of the 3D reconstruction of the LINC lattice (Fig. 4A, B).

**Fig. S5.**
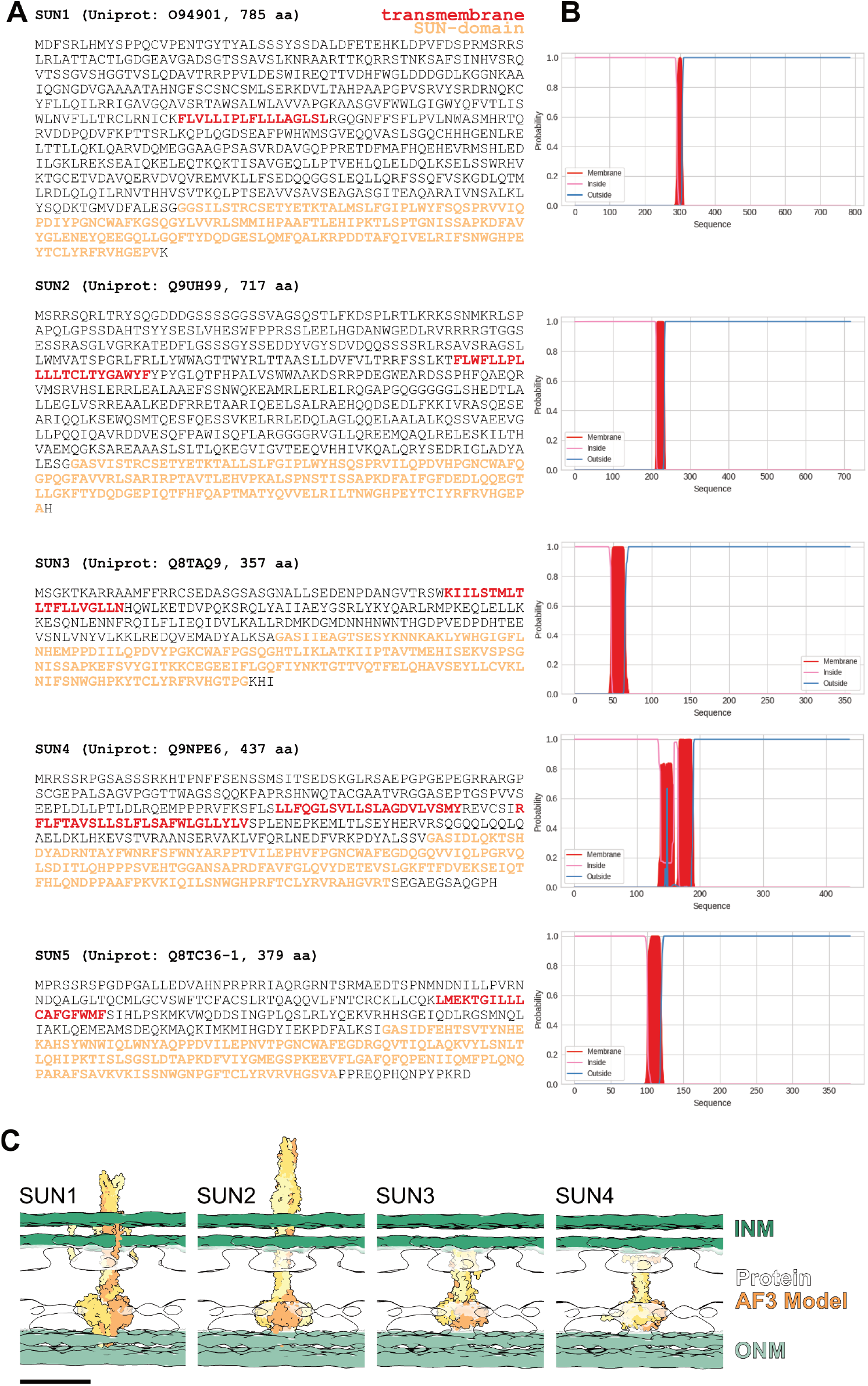
Prediction of human SUN protein transmembrane helices and their PNS-localized trimeric structures. **(A)** Sequences of human SUN proteins. Transmembrane regions predicted by DeepTMHMM (78) and SUN domains (according to corresponding UniProt entries) are highlighted. **(B)** DeepTMHMM plots displaying the probability of transmembrane helices being present in the corresponding SUN protein amino acid sequences. **(C)** Fitting of AF3 predictions of all human SUN proteins (SUN domain and coiled-coil) into the subtomogram average shown in Fig. 4 A-B. Scale bar, 10 nm.

**Fig. S6.**
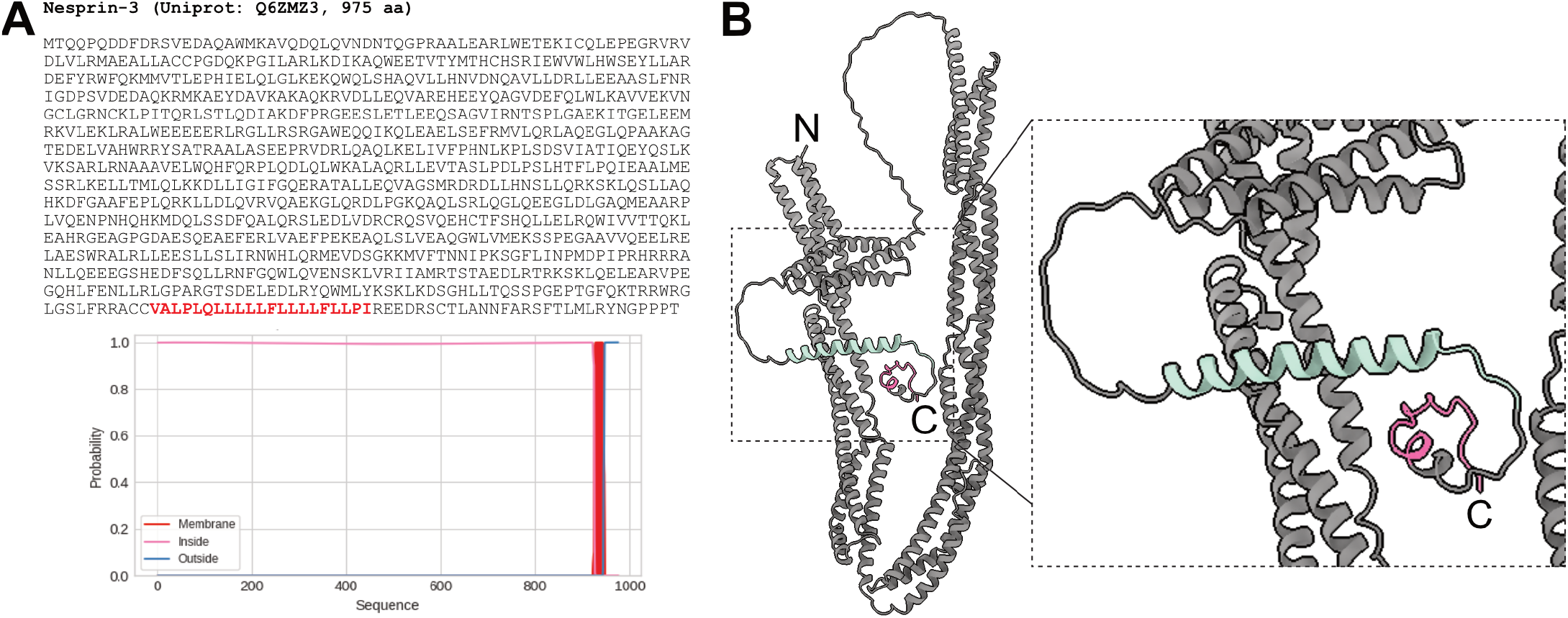
Transmembrane helix and AlphaFold3 structure prediction of Nesprin3. **(A)** Sequence and DeepTMHMM prediction of Nesprin3 transmembrane regions. **(B)** Structure prediction of Nesprin3 monomer. The SUN5 binding C-terminus is shown in pink and the predicted transmembrane helix is shown in green.

**Table 1.**
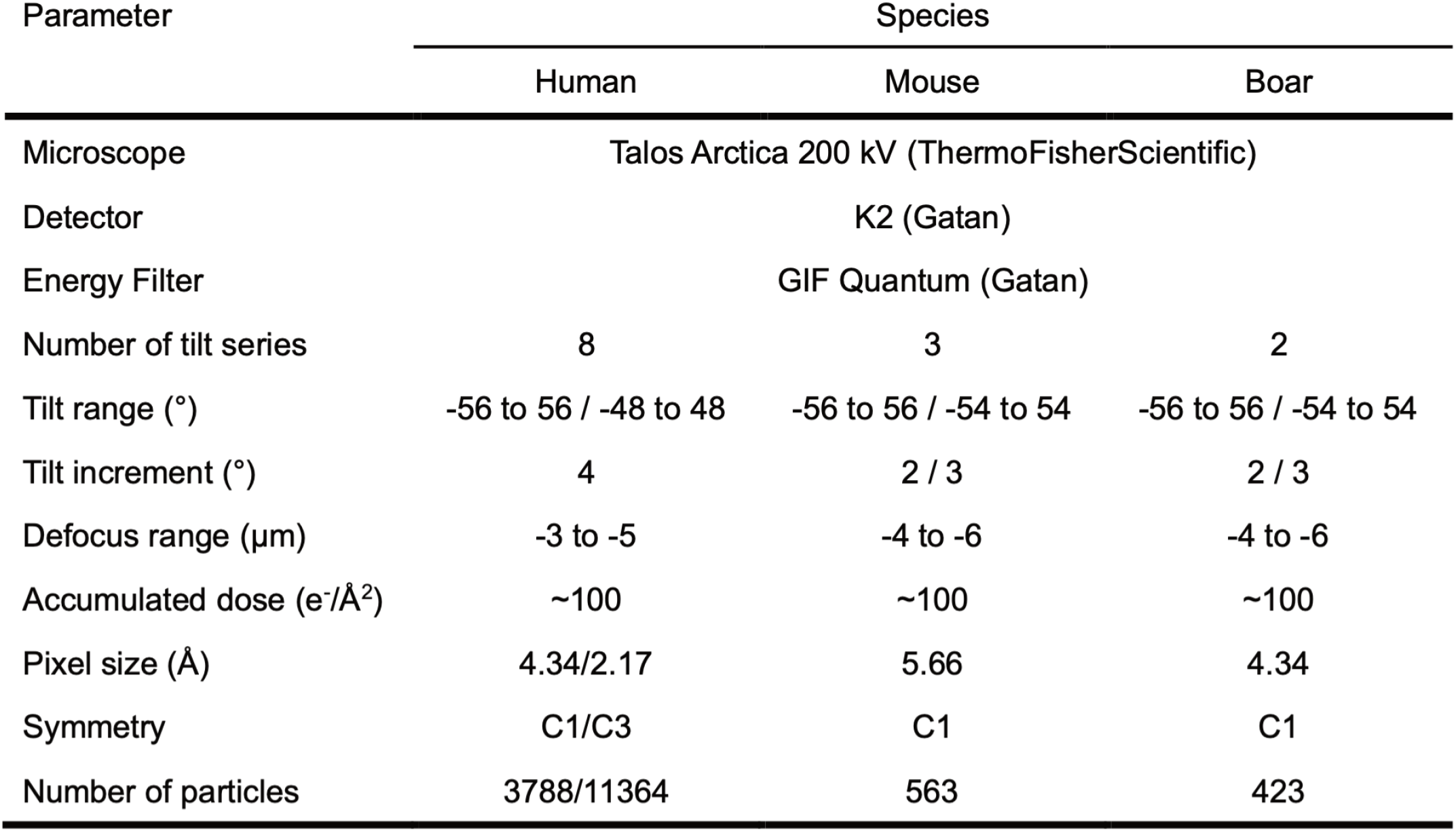
Subtomogram averaging summary.

| Parameter | Species |  |  |
| --- | --- | --- | --- |
|  | Human | Mouse | Boar |
| Microscope | Talos Arctica 200 kV (ThermoFisherScientific) |  |  |
| Detector | K2 (Gatan) |  |  |
| Energy Filter | GIF Quantum (Gatan) |  |  |
| Number of tilt series | 8 | 3 | 2 |
| Tilt range (°) | -56 to 56 / -48 to 48 | -56 to 56 / -54 to 54 | -56 to 56 / -54 to 54 |
| Tilt increment (°) | 4 | 2 / 3 | 2 / 3 |
| Defocus range (μm) | -3 to -5 | -4 to -6 | -4 to -6 |
| Accumulated dose (e<sup>-</sup>/Å<sup>2</sup>) | ~100 | ~100 | ~100 |
| Pixel size (Å) | 4.34/2.17 | 5.66 | 4.34 |
| Symmetry | C1/C3 | C1 | C1 |
| Number of particles | 3788/11364 | 563 | 423 |

